# New insights into the cultivability of human milk microbiota from ingestion to digestion and implications for its immunomodulatory properties

**DOI:** 10.1101/2024.12.11.626747

**Authors:** Charles Le Bras, Alizé Mouchard, Lucie Rault, Marie-Françoise Cochet, Olivia Ménard, Nolwenn Jacquet, Victoria Chuat, Florence Valence, Yves Le Loir, Amandine Bellanger, Amélie Deglaire, Isabelle Le Huërou-Luron, Sergine Even

**Affiliations:** STLO, INRAE, Institut Agro Rennes Angers, Rennes, France; Institut NuMeCan, INRAE, INSERM, Université de Rennes, Saint Gilles, France; CHU Rennes, Pediatric Department, CIC-Inserm 1414, Rennes, France

**Keywords:** Breast milk microbiota, gastrointestinal digestion, bacterial cultivability, bacterial viability, immunomodulatory properties, in vitro digestion

## Abstract

Human milk (HM) microbiota is increasingly studied for its potential health benefits. However, the physiological state of HM bacteria and consequently their effects on gut homeostasis remain a question. This study investigated the physiological state of the HM microbiota by characterizing its cultivable fraction, as it might be at the point of ingestion and assessing the effects of digestion on the cultivability and immunomodulatory properties of six prevalent HM strains. The microbiota of 28 HM samples was analysed by 16S metabarcoding either directly on raw milk (raw milk microbiota, RM) or on the complete cultivable fraction obtained from seven non-selective media (cultivable milk microbiota, CM). Diversity was lower in CM than in RM, with 32 versus 435 genera and a median of 7 versus 69 genera per sample in CM and RM, respectively. CM also showed under-representation of strictly anaerobic genera. Factors like parity and iron or vitamin supplementation affected RM and/or CM. *In vitro* gastrointestinal digestion moderately impacted strain cultivability. However, most strains partially or completely lost their immunomodulatory properties on the monocyte THP1 cell line after digestion, except *a Staphylococcus epidermidis* strain that gained immunomodulatory potential.

## Introduction

Human milk (HM) microbiota has been widely explored in recent years, mainly through metagenomic approaches, revealing an initially unsuspected diversity ^1–6^. Hence, HM was reported to host a low bacterial load but complex microbial community, with hundreds of bacterial genera and species identified, mainly within the Bacillota, Actinomycetota and Pseudomonadota ^2,7^. The most prevalent bacterial species include *Staphylococcus* and *Streptococcus*, that are universally predominant in HM as well as *Cutibacterium, Corynebacterium, Bifidobacterium, Lactobacillus, Pseudomonas, Rothia, Acinetobacter, Enterococcus* and several genera known to be strictly anaerobic such as *Fecalibacterium, Bacteroides* or *Veillonella* ^2–4,7^. A few studies have also investigated this microbial community using culture-dependent approaches, revealing a less diverse bacterial community than with culture-independent approaches, and a limited number of strictly anaerobic bacteria ^1,8–11^.

These culture-dependent approaches are generally based on the selection of a subset of isolates that are individually identified by sequencing the 16S rRNA gene or by mass spectrometry analysis^1,11^.

Discrepancies between culture-dependent and independent methods question our ability to capture the microbial diversity of HM with culture-dependent methods, but they also question the physiological state of HM bacteria and, consequently, their ability to colonize the infant gut and influence the infant gut microbiota and homeostasis through direct interactions and or their metabolic activities ^12,13^. Nevertheless, several studies support a role of HM microbiota on infant gut homeostasis. Some studies pointed to an overlap between the microbiota of HM and that of the infant gut or, more broadly, to the influence of the first one on the composition of the second one ^14–16^. In addition, some strains are shared between HM and the infant microbiota within a dyad, notably strictly anaerobic bacteria ^17,18^. HM microbiota is thought to play a wider role in intestinal homeostasis, immune and barrier functions ^19^. *In vitro* characterization of *Bifidobacterium*, *Lactobacillus* and *Streptococcus* species present in HM has highlighted their ability to modulate the immune and barrier functions of epithelial cells ^20–23^. Our previous work on a large number of HM strains has demonstrated the functional diversity of these bacteria in a quadricellular model of intestinal epithelium, both in terms of their ability to stimulate the model’s pro-and anti-inflammatory responses, and their capacity to reinforce or diminish the intestinal epithelial barrier ^24^. The question of the physiological status of HM bacteria goes beyond the state at the time of ingestion, as this state can be modified during digestion. *In vitro* characterization of HM strains is generally carried out using live bacteria derived from fresh cultures, which in all cases have not undergone digestion steps ^24^. The ability of bacterial strains to survive in the gastrointestinal tract has been mainly investigated for probiotic strains, especially *Bifidobacterium* and *Lactobacillus* strains ^25^. However, strong inter-strain variability has been reported with important loss of cultivability, up to several log ^26–28^. Survival to the digestion process remains rather unexplored in the context of HM which hosts a large bacterial diversity, beyond lactic acid bacteria and bifidobacteria. Furthermore, the digestive tract of newborns is immature and the digestion process differs from adults, with less harsh conditions in infants ^29^. Whether these milder digestive conditions in infants may allow a better survival of a large set of HM bacteria with consequences on their ability to interact with the intestinal epithelium remains to be evaluated.

In this study, we aimed to address the physiological state of HM microbiota by exploring its cultivable fraction at the time milk is expressed (i.e. corresponding to the point of ingestion by infant).

Cultivability has been used here as a proxy to estimate the viable fraction of the HM microbiota, albeit imperfectly, as it excludes the so-called “viable but non-cultivable” fraction. The total cultivable fraction of 28 HM was recovered from seven non-selective media and sequenced by 16S rRNA metabarcoding. This complete cultivable HM microbiota was compared to the raw milk microbiota obtained by 16S rRNA metabarcoding directly on HM samples. In addition to considering cultivability at ingestion, we evaluated the effect of the gastrointestinal digestion process, using an infant *in vitro* static digestion model, on the cultivability and immunomodulatory properties on the monocyte THP1 cell line, of six different HM strains.

## Results

### A lower diversity in the cultivable milk microbiota as compared to the raw milk microbiota

The first objective of this study was to assess the complete cultivable milk microbiota (CM) of 28 HM samples (see Supplementary Table S1 for sample description and associated metadata). For CM, each HM sample was plated on seven non-selective media under aerobic and/or anaerobic conditions in order to promote the growth of the greatest diversity of bacteria. This included a broad-spectrum rich medium, three media known to promote the growth of strictly anaerobic bacteria and two media that promote the growth of fastidious microorganisms and lactic acid bacteria. Of note, incubation was performed at 37°C for one to three days, which may limit the recovery of microorganisms with different optimal growth temperatures and/or low growth rates. For each HM sample, the cultivable fraction obtained under the seven growth conditions was collected by scraping the plates and pooling them before sequencing using 16S metabarcoding. This approach was used in order to obtain a complete overview of the CM of each HM sample, which may not be fully achieved by selecting and identifying a few isolates from each medium as usually done ^1,11^. CM was then compared with the raw milk microbiota (RM) obtained by direct metabarcoding on the 28 HM samples. Overall, our sequencing effort produced a total of ∼ 3.3 million quality filtered sequences corresponding to 27 RM and 28 CM (the available volume of one milk sample was too small to allow RM analysis). The median number of sequences was 53 465 sequences per sample (24772 and 91 741 in RM and CM, respectively), corresponding to 1775 bacterial OTUs detected based on a minimum abundance of >0.005% in the full dataset. The abundance table for each sample is presented in Supplementary Table S2 online, following aggregation at different taxonomic levels.

A general overview of RM and CM revealed that they strongly differed in terms of both α- and β- diversities (Figure 1). Hence, the α-diversity of CM was significantly lower than that of RM (Figure 1a, Table 1). Similarly, the sample type affected the β-diversity, with a clear separation between RM and CM samples (Figure 1b), and a contribution of sample type to beta-diversity between 17% and 53% (Table 2). The complete RM included 27 phyla corresponding to 435 genera, 605 species and 1231 OTUs whereas 4 phyla, 32 genera, 68 species and 612 OTUs were found in CM (Table 3). Likewise, the median number of genera and OTUs per sample was ∼10-fold and 2-fold lower in CM compared to RM (69 genera and 96 OTUs/samples for RM and 7 genera and 53 OTUs/sample for CM).

**Figure 1.**
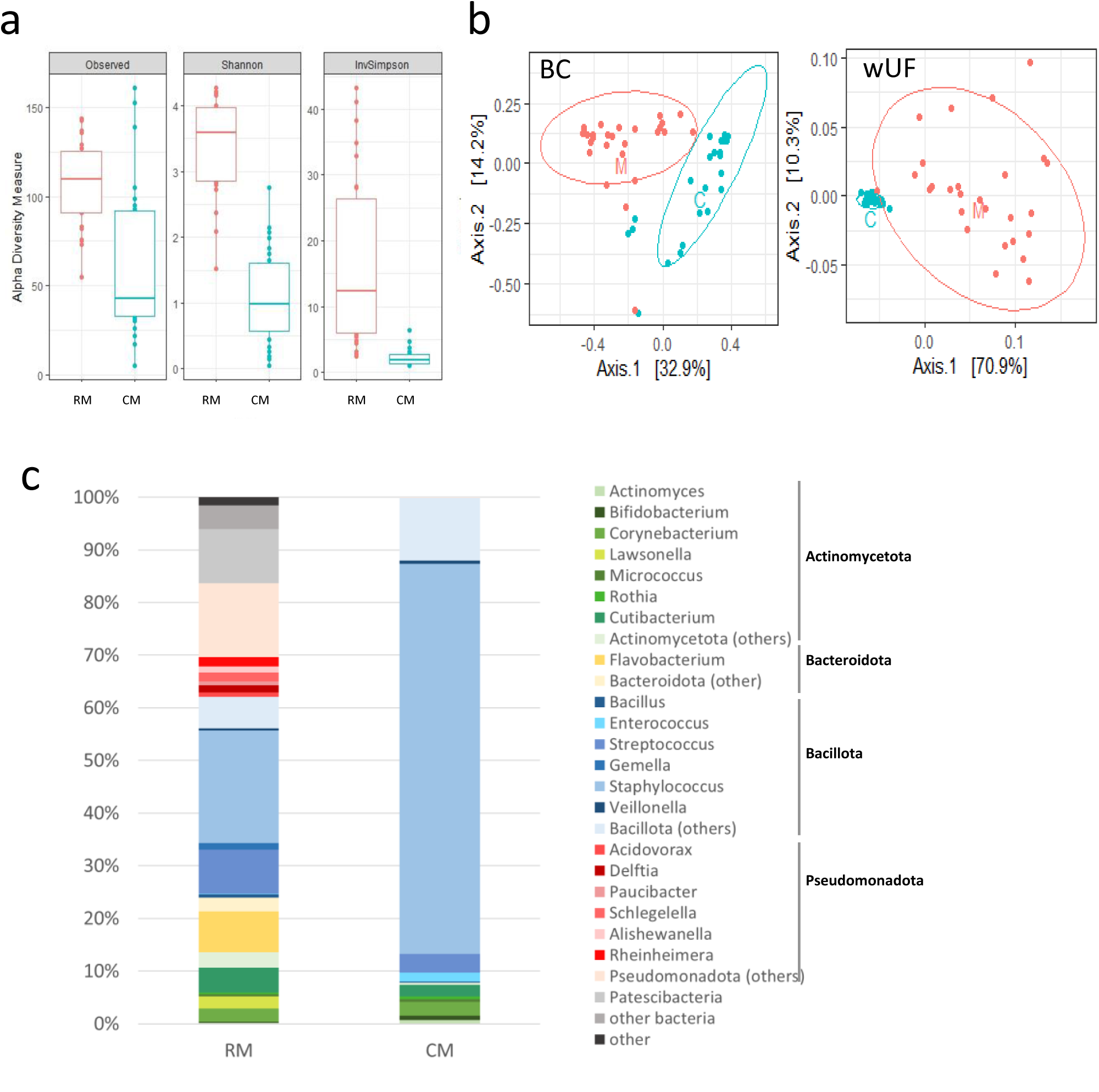
Overview of HM microbiota with regard to the sample type. **a.** Alpha-diversity (Observed richness, Shannon and Inverse Simpson index) of raw milk microbiota (RM) (red) and cultivable milk microbiota (CM) (blue). **b.** Multi-Dimensional Scaling (MDS) of RM and CM. MDS was performed based on the measurement of the Bray–Curtis (BC) or weighted UniFrac (wUF) distances. Samples are indicated by points and coloured with regard to the sample type: RM (red) and CM (blue). **c.** Mean taxonomic profiles of RM and CM. The 20 dominant genera are presented as well as the 5 dominant phyla. Genera belonging to Actinomycetota (formerly named Actinobacteria) are displayed in shades of green, Bacteroidota (formerly named Bacteroidetes) in shades of yellow, Bacillota (formerly named Firmicutes) in shades of blue, Pseudomonadota (formerly named Proteobacteria) in shades of red.

**Table 1.** Effects of different factors on the α-diversity of raw and cultivable milk microbiota or both.

| Factor <sup>1</sup> | Observed |  | Chao |  | Shannon |  | Simpson |  | InvSimpson |  |
| --- | --- | --- | --- | --- | --- | --- | --- | --- | --- | --- |
| **raw and cultivable milk microbiota** |  |  |  |  |  |  |  |  |  |  |
| type | 2.65E-06 | *** | 0.007 | ** | 1.35E-16 | *** | 3.34E-12 | *** | 1.95E-08 | *** |
| parity | 0.039 | * | 0.016 | * | 0.971 |  | 0.326 |  | 0.021 | * |
| iron_G | 0.895 |  | 0.917 |  | 0.301 |  | 0.910 |  | 0.040 | * |
| vit_BF | 0.018 | * | 0.044 | * | 0.065 | . | 0.031 | * | 0.902 |  |
| **raw milk microbiota** |  |  |  |  |  |  |  |  |  |  |
| parity | 0.250 |  | 0.152 |  | 0.058 | . | 0.058 | . | 0.013 | * |
| iron_G | 0.164 |  | 0.148 |  | 0.033 | * | 0.033 | * | 0.021 | * |
| vit_BF | 0.612 |  | 0.727 |  | 0.994 |  | 0.994 |  | 0.905 |  |
| **cultivable milk microbiota** |  |  |  |  |  |  |  |  |  |  |
| parity | 0.097 | . | 0.082 | . | 0.042 | * | 0.076 | . | 0.095 | . |
| iron_G | 0.495 |  | 0.443 |  | 0.711 |  | 0.874 |  | 0.778 |  |
| vit_BF | 0.017 | * | 0.022 | * | 0.004 | ** | 0.016 | * | 0.0013 | ** |

<sup>1</sup> Summary of the effects (P values) of different factors on different α-diversity indexes (Observed richness, Chao, Shannon, Simpson and Inverse Simpson index) of the raw milk microbiota (RM) and cultivable milk microbiota (CM) as determined by ANOVA with several factors: type, either RM or CM; parity: either primiparous or multiparous; iron\_G: mothers supplemented or not by iron during gestation; vit\_BF : mothers supplemented or not by vitamins during breastfeeding. \*\*\*P value < 0.001; \*\*P value < 0.01; \*P value < 0.05; "." P value < 0.1

**Table 2.**
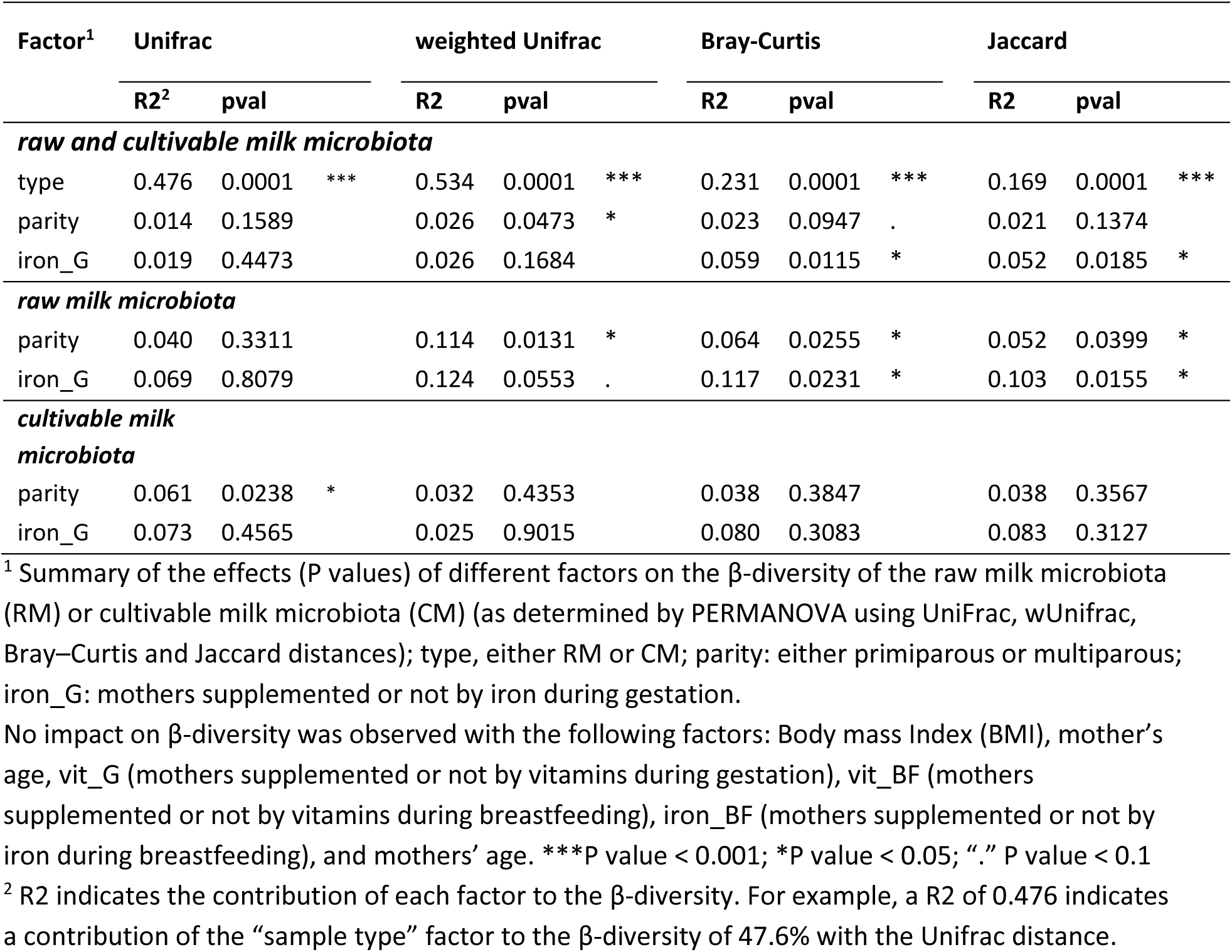
Effects of different factors on the β-diversity of raw and cultivable milk microbiota or both.

**Table 3.**
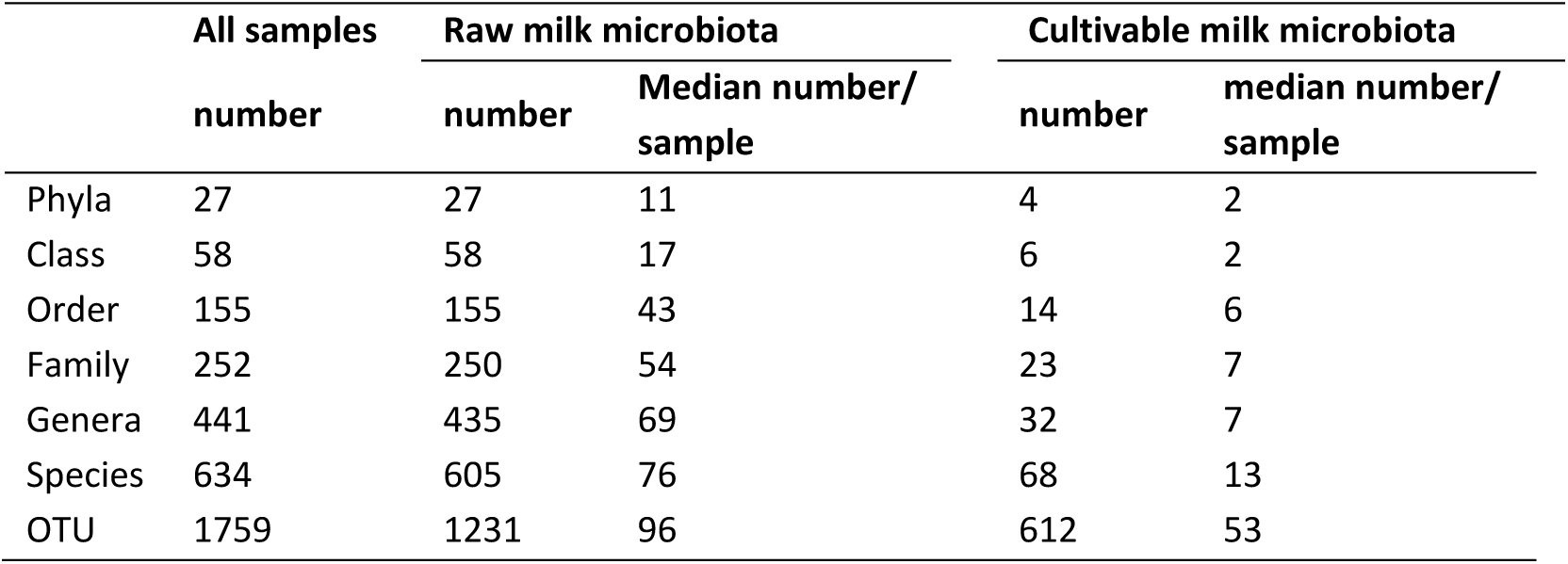
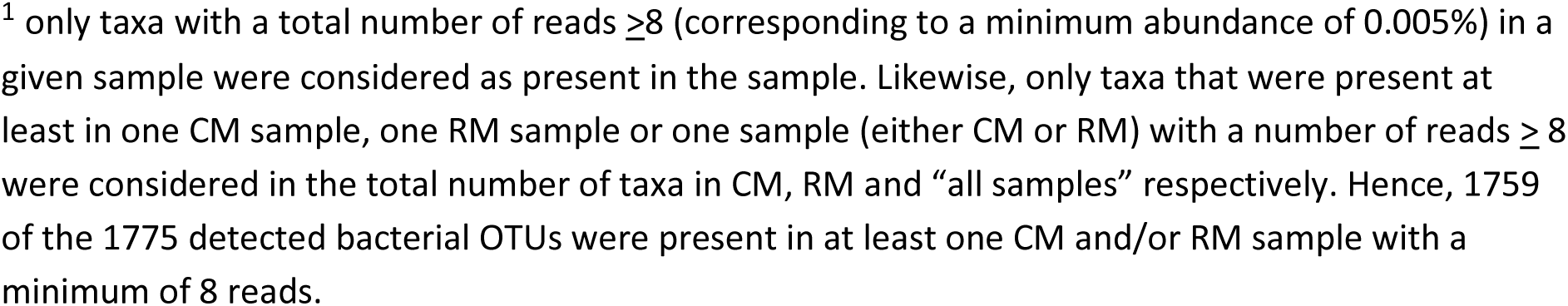
Overview of the bacterial taxonomic composition of raw milk microbiota (RM) and cultivable milk microbiota (CM). Total number of taxa for each taxonomic level is given as well as the median number per sample^1^.

|  | All samples | Raw milk microbiota |  | Cultivable milk microbiota |  |
| --- | --- | --- | --- | --- | --- |
|  | number | number | Median number/<br>sample | number | median number/<br>sample |
| Phyla | 27 | 27 | 11 | 4 | 2 |
| Class | 58 | 58 | 17 | 6 | 2 |
| Order | 155 | 155 | 43 | 14 | 6 |
| Family | 252 | 250 | 54 | 23 | 7 |
| Genera | 441 | 435 | 69 | 32 | 7 |
| Species | 634 | 605 | 76 | 68 | 13 |
| OTU | 1759 | 1231 | 96 | 612 | 53 |

<sup>1</sup> only taxa with a total number of reads $\geq 8$ (corresponding to a minimum abundance of 0.005%) in a given sample were considered as present in the sample. Likewise, only taxa that were present at least in one CM sample, one RM sample or one sample (either CM or RM) with a number of reads $\geq 8$ were considered in the total number of taxa in CM, RM and “all samples” respectively. Hence, 1759 of the 1775 detected bacterial OTUs were present in at least one CM and/or RM sample with a minimum of 8 reads.

RM was dominated by 5 phyla, namely Bacillota (mean abundance of 38%), Actinomycetota (14%), Pseudomonadota (21%), Bacteroidota (10%) and Patescibacteria (10%), while CM was almost exclusively composed of the two formers with mean relative abundance of 92% and 8% (Figure 1c). Bacteroidota and Pseudomonadota were almost absent in CM at the exception of *Prevotella*, *Moraxella, Klebsiella* and *Haemophilus*. *Staphylococcus*, *Streptococcus, Cutibacterium* and *Corynebacterium* were the most abundant and prevalent genera in CM. Additional prevalent and/or abundant genera present in CM included *Enterococcus, Bacillus, Veillonella, Bifidobacterium, Micrococcus, Kocuria, Actinomyces* and *Rothia*, and, to a lesser extent *Lactobacillus, Finegoldia* and *Granulicatella* (Supplementary Figure S1 and Table S2). While all the genera present in CM were present in RM as well, several genera present in RM were not recovered from CM. This included strictly anaerobic genera such as those belonging to the Clostridia and Negativicutes classes, with the exception of *Finegoldia* and *Veillonella*, which were present in CM of few samples (Supplementary Table S2).

### Factors influencing human milk microbiota richness and composition

Several metadata were associated with the HM collection, allowing to evaluate the impact of several factors on HM microbiota (Tables 1 and 2). Parity was found to affect both RM and CM α-diversity, yet in opposite directions. In RM, milk from primiparous mothers had a lower α-diversity than that of multiparous, as illustrated with the Inverse Simpson index (and a trend with Shannon and Simpson indexes). Conversely, in CM, α-diversity was higher in primiparous compared to multiparous mothers, as illustrated on the Shannon index (and a trend with other indexes) (Table 1, Supplementary Figure S2). Parity also contributed to RM β-diversity with a contribution between 4 and 11%, and, to a lesser extent, CM β-diversity (Table 2). Several taxa showed differential abundance in RM and CM from primiparous compared to multiparous mothers (Supplementary Tables S3 and S4).

Supplementations during the gestation or breastfeeding also influenced α- and β-diversities. Mother supplemented with iron during the gestation or with vitamins during the breastfeeding period had a higher α-diversity in RM and CM respectively (Table 1, Supplementary Figure S2). Iron supplementation during the gestation also influenced RM β-diversity, with a contribution of 7 to 12% (Table 2). Differential abundance analysis showed that iron supplementation during the gestation and vitamin supplementation during breastfeeding modulated the abundance of several taxa (Supplementary Tables S3 and S4).

### Infant gastrointestinal digestion affects bacterial cultivability and immunomodulatory properties in a strain dependant manner

A second objective was to evaluate the impact of the gastrointestinal digestion on the cultivability and immunomodulatory properties of 6 strains previously isolated from HM and belonging to prevalent species of HM microbiota: *Bifidobacterium breve* CIRM BIA 2845, *Streptococcus salivarius* CIRM BIA 2846, *Cutibacterium acnes* CIRM BIA 2849, *Staphylococcus epidermidis* CIRM BP 1633, *Lactobacillus gasseri* CIRM BIA 2841 and *Enterococcus faecalis* CIRM BIA 2835 ^24^. The six strains were tested using an *in vitro* digestion model adapted to the infant stage ^29^. The gastric phase of digestion had no significant impact on bacterial population, with the exception of *E. faecalis* whose population slightly increased (0.3-log) (Figure 2). On the contrary, the intestinal phase impacted the bacterial cultivability in a strain-dependant manner, with a decrease of population of 2-log, 4.5-log and 0.3-log for *B. breve*, *S. salivarius* and *L. gasseri* respectively, a 0.7-log increase for *E. faecalis* compared to the initial population and no impact for *C. acnes* and *S. epidermidis* (Figure 2). All strains retained at least partial cultivability after the gastrointestinal digestion.

**Figure 2.**
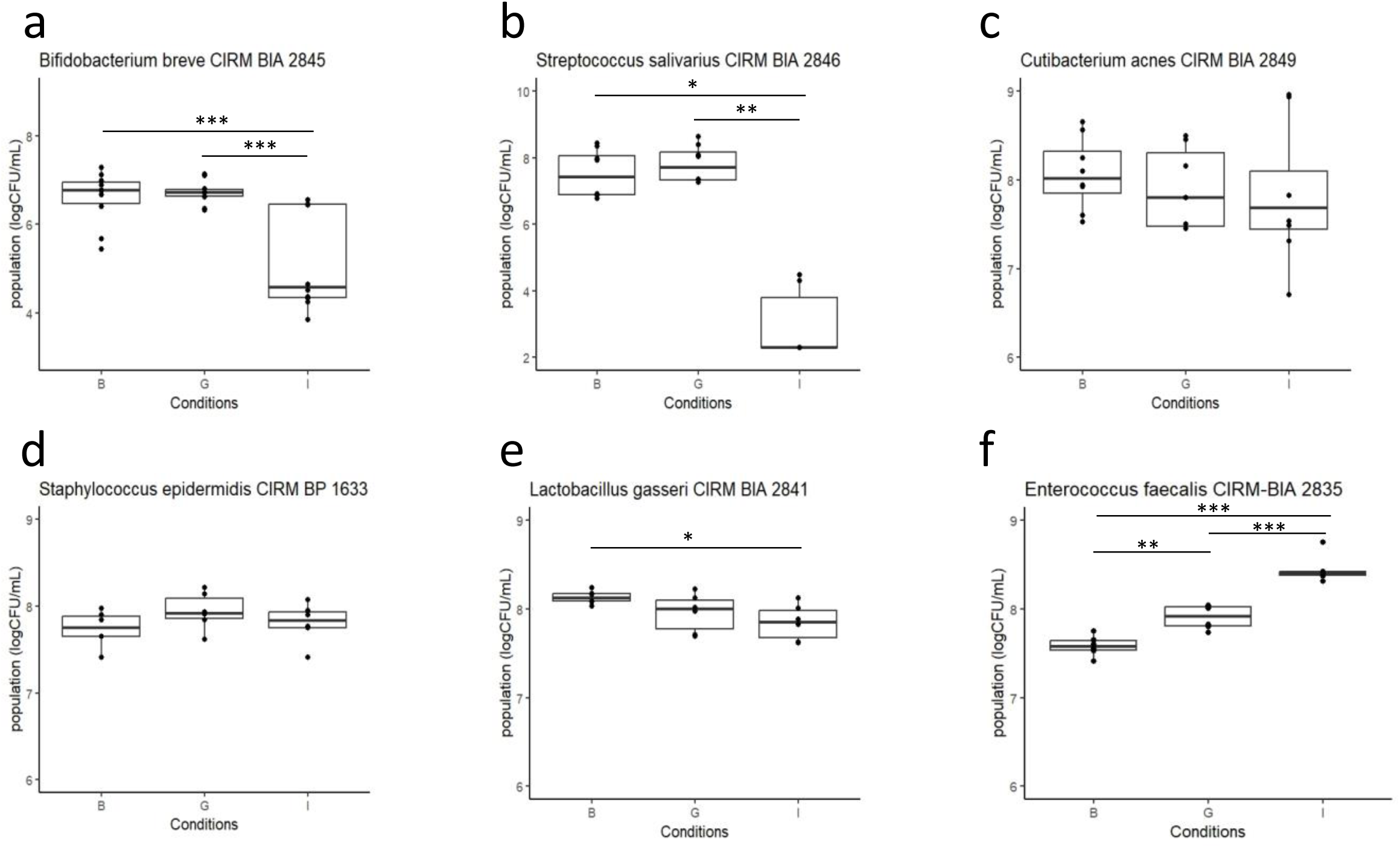
Impact of gastric (G) and gastro-intestinal (I) digestion on the cultivability of six HM strains, as compared to the control before digestion (B). Strains were added to an infant formula and digested under static conditions mimicking the infant gastric (G) or gastric followed by intestinal digestion (I). Cultivable population of each strain was determined on its optimal growth medium. Significant changes in the cultivable population were determined by ANOVA followed by a post-hoc test to find significant pairwise differences. ***P value < 0.001; **P value < 0.01; *P value < 0.05. a; *Bifidobacterium breve* CIRM BIA 2845; b: *Streptococcus salivarius* CIRM BIA 2846; c: *Cutibacterium acnes* CIRM BIA 2849; d: *Staphylococcus epidermidis* CIRM BP 1633; e: *Lactobacillus gasseri* CIRM BIA 2841; f: *Enterococcus faecalis* CIRM BIA 2835.

We further investigated the effect of digestion on the immunomodulatory properties of the 6 strains on THP1 cells differentiated into macrophages, through the production of IL-10 and TNF-α. We used two different controls, cells without bacterial stimulation (B) and cells stimulated with the digested formula alone without bacteria (IF-I). The digested infant formula had no effect on TNF-α release by THP1 cells, but induced a slight decrease in IL-10 release (Figure 3). The immunomodulatory properties were modulated by the digestion process in a strain-dependent manner. *C. acnes and L. gasseri* completely lost their immunomodulatory properties, as shown by the absence of differences in IL10 and TNF-α production between cells stimulated with either the digested strain (I) or the digested IF alone (IF-I). (Figure 3e,f,i,j). After gastrointestinal digestion, *E. faecalis* was still able to slightly stimulate TNF-α secretion but had no more effect on IL-10 production compared to the digested formula alone (Figure 3k,l). Conversely, S*. salivarius* still very slightly but significantly decreased IL-10 production compared to the digested formula alone but had no more effect on TNF- α secretion (Figure 3c,d). *B. breve* partially retained its ability to stimulate IL-10 and TNF-α compared to the digested formula alone (Figure 3a,b). Finally, *S. epidermidis* gained the ability to slightly stimulate IL-10 and increased its stimulation of TNF-α production (Figure 3g,h).

**Figure 3.**
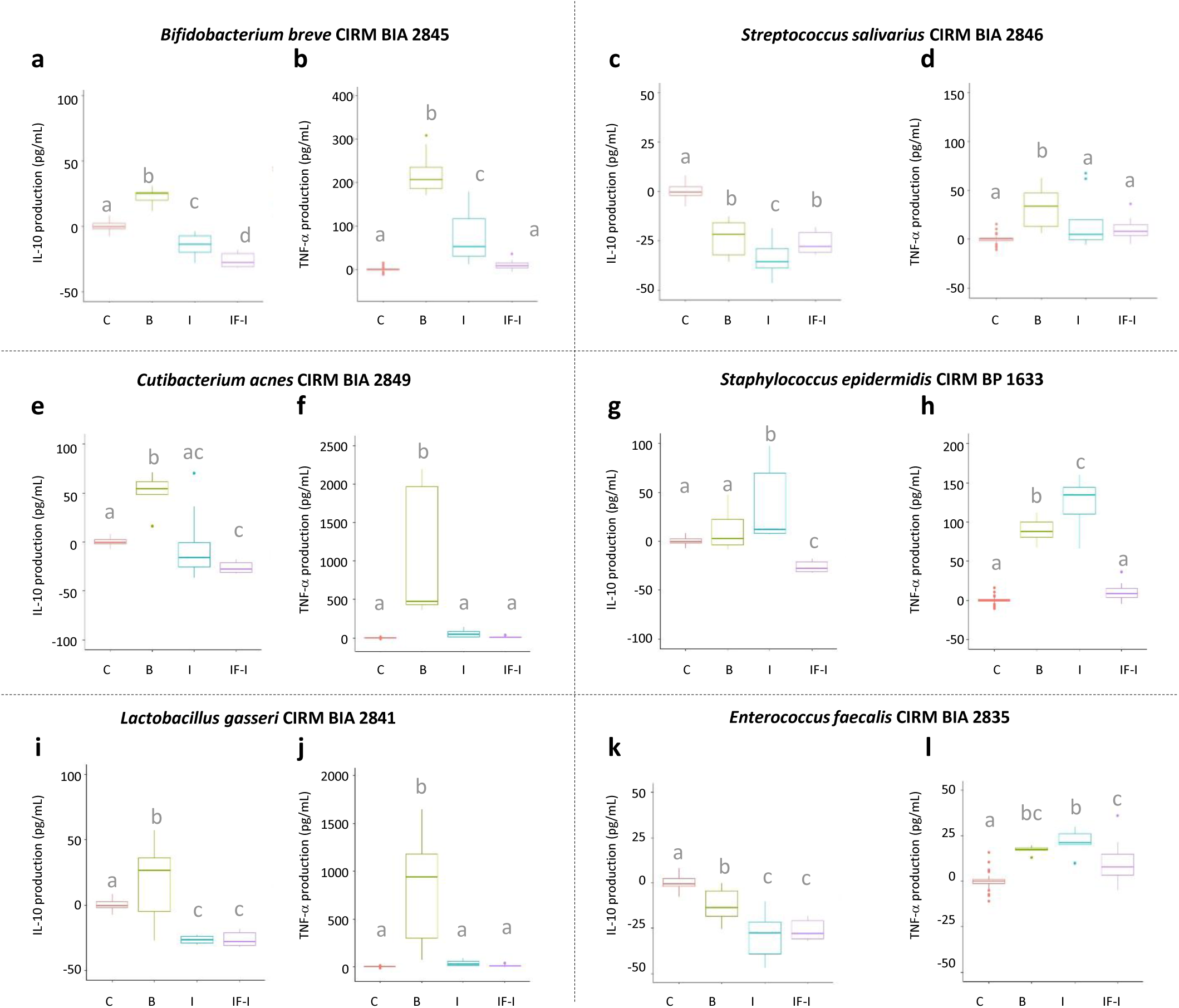
Impact of gastro-intestinal digestion on the immunomodulatory properties of six HM strains. Strains were digested in infant formula under static conditions mimicking the neonate gastric followed by intestinal digestion, recovered by centrifugation before assessing their immunomodulatory properties on THP1 cells through the impact on IL-10 (a, c, e, g, I, k) and TNF-α (b, d, f, h, j, l) secretion. Bacteria, either before (B) and after gastro-intestinal digestion (I), were added at a MOI of 10 bacteria per cell. IL-10 and TNF-α production (pg/mL) were corrected by the production in the absence of bacterial stimulation (C, control) of the corresponding replicate. An additional control corresponded to the digestion of the infant formula without the addition of bacteria (IF-I). Significant changes in IL-10 or TNF-α production were determined by ANOVA followed by a post-hoc test to find significant pairwise differences. Different letters correspond to conditions that were significantly different (P value < 0.05). a, b; *Bifidobacterium breve* CIRM BIA 2845; c, d: *Streptococcus salivarius* CIRM BIA 2846; e, f: *Cutibacterium acnes* CIRM BIA 2849; g, h: *Staphylococcus epidermidis* CIRM BP 1633; i, j: *Lactobacillus gasseri* CIRM BIA 2841; k, l: *Enterococcus faecalis* CIRM BIA 2835

## Discussion

Although long ignored, the milk microbiota is now considered to be a component of HM that may contribute to infant gut microbiota and homeostasis ^12,13^. Despite numerous studies on HM microbiota, the question of the physiological state of HM bacteria, that may influence gut colonisation and interactions with the host, remains largely unanswered. In this study, we addressed this issue from ingestion to the end of the small intestine, by exploring the complete cultivable fraction of fresh HM and assessing the effect of gastrointestinal digestion on the cultivability and immunomodulatory properties of 6 prevalent HM microbiota species. The cultivable HM microbiota is a subset of the viable HM microbiota. Some viable microorganisms of the HM may not be recovered under the growth conditions used and constitute the “viable but non-cultivable” fraction. Bacterial viability is a complex issue that encompasses a continuum of physiological states related to bacterial cell integrity, the ability of strains to grow (under the optimal laboratory growth conditions or the actual conditions) or to be metabolically active. While the ability to grow under actual gut conditions is a prerequisite for gut colonisation, non-growing and non-living bacterial cells may still interact with the host, to some extent, as reported for heat-killed probiotic strains ^30^.

The assessment of the cultivable HM microbiota by metabarcoding, as performed in the present study, led to the identification of 32 genera and 68 species in total, with a median number of 7 genera and 13 species per HM sample. In a related study, based on the same HM samples and aimed at establishing and characterising a collection of HM isolates, we individually identified a subset of 1245 isolates by 16sRNA sequencing. These isolates (∼5 isolates per HM sample per growth condition) were obtained on the same growth conditions as those used in the present study, plus two additional conditions (MRSc and WCc supplemented with mupirocin in order to inhibit the growth of the dominant Staphylococci and favour the isolation of additional taxa). These 1245 isolates corresponded to a total of 26 different genera and 59 species, with a median number of 5 genera and 7 species per HM sample ^24^. Thus, the complete assessment of CM by metabarcoding as performed in the present study, resulted in a moderate overall gain in diversity compared to the more conventional approach of individually identifying a subset of isolates^24^, especially when considering the bacterial diversity per sample (∼twice the number of species per sample). Of note, a moderate increase (∼8.5%) in the diversity of fecal microbiota has previously been reported using culturomics when picking and identifying all colonies on the plate compared to “experienced” colony picking (i.e., picking 2-3 colonies per colony type on each plate) ^31^. Our results on the diversity of CM are in the same range as the total diversity of CM reported by Treven et al., who combined culture-based methods with MALDI-TOF mass spectrometry identification and recovered 25 genera from 1086 isolates from 31 HM samples ^11^. Using MALDI-TOF mass spectrometry, Wang et al identified 14 genera and 34 species from >3500 colonies from 9 fresh HM samples ^32^.

Despite the more complete assessment of the cultivable fraction of HM by metabarcoding, which allowed some gain in diversity in each sample, our results confirm the low bacterial diversity in CM compared to RM. A total of 435 genera were recovered in RM from the 28 HM samples, with a median of 69 genera per sample. This high bacterial richness in HM has been widely documented and the main taxa identified here were in good agreement with the literature ^2–5,7^. However, several taxa present in the RM were not recovered from the CM, as illustrated by the ∼14-fold lower total number of genera and the ∼10-fold lower median number of genera per sample in CM compared to RM. CM was mainly composed of Bacillota and Actinomycetota, with *Staphylococcus, Streptococcus, Cutibacterium* and *Corynebacterium* being the most abundant and prevalent genera in CM, in agreement with previous studies ^1,11,32,33^. Other prevalent genera in CM included *Bacillus, Granulicatella, Bifidobacterium, Enterococcus, Veillonella* as well as *Lactobacillus, Kocuria, Micrococcus, Rothia, Finegoldia,* and *Actinomyces,* which have also been reported in the cultivable fraction of milk ^7,32^. It should be noted that the prevalence of genera was generally consistent between RM and CM or higher in RM than CM (Supplementary Figure 1), as previously reported ^11^, suggesting that some species or strains within the genera commonly found in RM and CM, were not cultivable under our growth conditions.

A major difference between RM and CM was the absence in CM of several strictly anaerobic genera, belonging to the phyla Pseudomonadota and Bacteroidota or to the Classes Clostridia and Negativicutes within Bacillota, although we used several media known to promote the growth of anaerobic bacteria ^10^. In addition, precautions were taken to preserve this anaerobic flora, as milk samples collected by the mothers were stored in an anaerobic bag immediately after sampling. Even a brief exposure to oxygen during sample processing or breastfeeding may have been sufficient to kill these highly oxygen-sensitive bacteria. We also cannot rule out the possibility that they were already in a non-living or non-cultivable state in HM. Nevertheless, Schwab et al reported an improved isolation of obligate anaerobic species, including strictly anaerobic Bacteroidota or Clostridia, after milk storage for 6 days at 4◦C, suggesting that these taxa are part of the cultivable fraction of HM microbiota ^10^. Similarly, greater bacterial diversity, particularly within the anaerobes, was reported by Wang et al after enrichment of 9 fresh HM in blood culture medium for 3-6 days (21 genera and 54 species compared to 14 genera and 34 species identified in fresh milk) ^32^. High-throughput culture approaches could help to isolate this oxygen-sensitive flora, based on the culturomic approaches developed for gut microbiota and genomic analysis of the nutrient requirements of HM microbiota ^34^.

The second main objective of this study was to evaluate the ability of six strains belonging to prevalent HM genera to survive during the infant digestion process, which was achieved thanks to the use of a static digestion model specifically calibrated for infants ^29^. Of note, the strains were digested in an infant formula, to mimic HM to some extent, as the matrix has previously been shown to affect the strain survival during digestion ^26,28,35^. Cultivability of the six strains were hardly affected by the gastric phase of digestion but it was affected by the intestinal phase of digestion in a strain-dependant manner, with *S. salivarius* and *B. breve* being most affected, *L. gasseri*, *C. acnes* and *S. epidermidis* being little or not affected, and with a slight increase in *E. faecalis* population. Strain differences in survival to digestion process have been reported previously, mainly for lactic acid bacteria such as *Lactobacillus* and *Bifidobacterium* ^26–28,36^. The good survival of lactic acid bacteria in the digestive tract is related to the ability of strains to adapt to acid stress or counteract the effects of bile salts, especially in relation to the expression of bile salt hydrolases ^37–39^. Good survival and even growth was expected from *E. faecalis*, known as a commensal inhabitant of the human gut ^40^. More interestingly, this study highlighted that non-lactic acid bacteria and non-gut-associated bacteria such as *Cutibacterium acnes* and *Staphylococcus epidermidis* retained high cultivability under the harsh physiological conditions encountered in the stomach and small intestine related to low pH and secretion of bile salts. In line with this, good survival was previously reported for a strain of *Propionibacterium freudenreichii* in different dairy matrices, *Cutibacterium* and *Propionibacterium* being closely related taxonomically ^41^. Likewise, Adouard et al evaluated the cultivability of a mixture of six bacteria and three yeasts from the microflora of surface-ripened cheeses in a three-compartment dynamic gastrointestinal digester (DIDGI). Compared to the other bacteria, a *Staphylococcus equorum* strain, in pure culture, showed an intermediate resistance to the whole digestion process, with a survival rate lower than *Hafnia Alvei*, similar to that of *Corynebacterium casei* and *Brevibacterium aurantiacum*, and higher than *Lactococcus lactis* and *Arthrobacter arilaitensis* ^42^. It should be noted that among *Staphylococcus* species, *Staphylococcus aureus* has several acid resistance mechanisms ^43^, as well as a bile efflux pomp and a bile-induced oxidoreductase that may mediate cholate resistance and contribute to survival under human colonic conditions ^44^.

Although the digestion process affected the cultivability of some strains, all strains were still partially cultivable after gastrointestinal digestion, with populations between 3 and 8.5 Log CFU/mL. The digestive tract of infant is still immature, with a higher gastric pH and a lower bile salt concentration than in adults ^29^. Thus, due to the less stressful conditions in the infant digestive tract, most ingested bacteria are likely to retain some viability and, subsequently, have a greater impact on the gut homeostasis than in adults, by colonizing the gut environment and/or interacting with the gut epithelium. It should be noted that the 6 HM strains used here were grown under optimal laboratory growth conditions before being subjected to the digestion process, and the effect of digestion on these strains in freshly expressed HM could be different in relation to their physiological state under these real-life conditions. Another limitation of the static model used here is that it mimics the digestion process through two sequential stages (the gastric and intestinal phases) and does not take into account the dynamics of changes throughout the digestive tract, in terms of pH and digestive enzymes. The use of a dynamic model of digestion such as the DIDGI^45^ would allow a more reliable assessment of the effect of digestion on the cultivability of bacterial strains. Nevertheless, the changes in conditions experienced by the bacteria in this static model are more drastic and therefore likely to be more stressful than in a dynamic model or *in vivo* where they are more gradual, suggesting that most of the strains used in this study would retain partial cultivability under dynamic digestion.

Despite a limited impact on strain cultivability, a partial or complete loss of immunomodulatory properties was revealed for most strains after digestion, with the exception of *Staphylococcus epidermidis*, which gained the ability to stimulate IL-10 production, and showed increased stimulation of TNF-α production. A loss of its anti-inflammatory properties was also reported for *Propionibacterium freudenreichii* after digestion ^41^. This loss is likely due to proteolysis of bacterial cell surface proteins involved in cell stimulation by digestive enzymes such as trypsin. In our study, the loss of immunomodulatory properties was not necessarily associated with the cultivability after gastrointestinal digestion, as illustrated by *C. acnes*, which completely lost its immunomodulatory properties, while cultivability was unaffected (Figures 2 and 3). Nevertheless, good survival can help to counteract the effects of proteolysis and allow *de novo* biosynthesis of new cell wall proteins.

Moreover, bacteria still cultivable after gastrointestinal digestion could theoretically grow or at least be metabolically active in the lower part of the digestive tract, where conditions are “friendlier”. The differences between strains could be related to the nature of the determinants involved in the interaction with THP1 cells such as proteins but also exopolysaccharides, teichoic acids, peptidoglycans or metabolites ^46–48^. Regarding *S. epidermidis*, it is possible that its immunomodulatory effects are related to non-protein components, or that the digestion of some surface components (proteins, polysaccharides) has uncovered other cell-interacting components, thus modifying the interplay with some intestinal cell receptors. From a more general point of view, this study highlights the need to take into account the effect of the digestion process not only on the cultivability but also on the properties of bacterial strains, given the poor relationship between these two parameters.

Finally, several metadata were associated with the HM collection, allowing to investigate the impact of several factors on RM and CM. Parity affected both RM and CM α-diversity, but in opposite directions. RM α-diversity was higher in multiparous than in primiparous mothers while the reverse was true for CM. Although there is no consensus on the effect of parity ^7^, our results corroborate a previous study that reported higher α-diversity in HM from multiparous mothers ^49^. This apparent discrepancy between RM and CM suggests a selective enrichment of HM with non-cultivable bacteria between the first and subsequent lactations. The enrichment of RM with the number of lactations is probably a result of the bidirectional cross-talk between mother and infant through retrograde flow, allowing milk enrichment by infant oral and maternal skin microbiota ^49^. We also cannot exclude that the enrichment of multiparous RM with non-cultivable taxa is due to the colonization of the mammary gland by maternal gut bacteria during the first lactation via the entero-mammary pathway^33,50^.

Other factors affecting RM and CM include iron supplementation during gestation and vitamin supplementation during breastfeeding, which were associated with a higher α-diversity in RM and CM, respectively, and several changes in composition. While several studies have documented the relationship between HM microbiota and maternal diet, including protein and fibre intake or unsaturated fatty acids ^6,7,51^, the effect of vitamin or iron supplementation on HM microbiota is not or poorly documented. Relations were reported between *Streptococcus* abundance in the HM microbiota and folic acid or vitamin B-12 intake in the first months of lactation ^54^, and between the level of Vitamin C intake during gestation and the HM microbiota profile ^55^. Iron has also been shown to affect the richness and composition of the gut microbiota ^56^. Specifically, Pereira et al reported reduced gut microbiota richness in rats following dietary-induced iron-depletion, whereas it was partially restored following iron supplementation ^57^. Nitert et al found no differences in fecal microbiota richness in overweight women in early pregnancy receiving high or low dose iron supplementation, but a limited but significant effect on β-diversity was observed ^58^. In our study, the changes in milk microbiota composition induced by iron supplementation during gestation may have been induced by those in the gut microbiota through the entero-mammary pathway ^3,6,33^. Of note, due to the small sample size of our study, our findings on factors influencing milk microbiota, particularly iron and vitamin supplementation during gestation or lactation, should be considered with caution and confirmed by additional studies in larger cohorts.

In conclusion, the complete assessment of the HM cultivable microbiota by metabarcoding, as proposed in the present study, enabled a moderate gain in diversity within each HM sample compared to a more conventional approach based on the individual identification of a subset of isolates. Nevertheless, we confirmed that the diversity of the CM was limited compared to that of the RM. Furthermore, the gastrointestinal digestion process, as it occurs in the immature infant digestive tract, affected bacterial cultivability in a strain-dependent manner, with overall good maintenance of cultivability for most HM strains including non-lactic acid bacteria, or non-gut-associated bacteria. This suggests that most of them could grow and/or be metabolically active in the small intestine and later in the colon. Similarly, the immunomodulatory properties of HM bacteria were modulated in a strain-dependent manner, with no direct relationship to cultivability. Most of the strains tested partially or completely lost their immunomodulatory properties on THP1 cells, with the exception of a *Staphylococcus epidermidis* strain, which gained some immunomodulatory capacity. These results highlight the potential of HM bacteria, regardless of their taxonomy, to interact with the intestinal immune system, and the importance of considering the digestion process when addressing the question of their interactions with the intestinal epithelium and mucosal immune system.

## Material and Methods

### HM sampling and preparation of the cultivable HM fraction

HM sampling has been described previously ^24^. Briefly, the HM sampling protocol was approved by the Institutional Review Board of the Poitiers Hospital (n°20.05.27.67526). HM sampling was performed between 2.0 and 6.0 weeks post-partum as previously described ^24^, on twenty-eight healthy mothers who delivered a healthy baby vaginally at term and exclusively breastfed their infant. Exclusion criteria were any formula feeding in addition to breast-feeding, signs of infection or administration of drugs (including antibiotics) to the baby and to the mother during the 3 months prior to delivery and during lactation. HM samples were stored at-80°C for direct analysis of the raw milk microbiota (RM). In addition, each fresh HM sample was used for the determination of the cultivable milk microbiota (CM). HM samples (100 µL) were plated on seven different non-selective media to promote the growth of the greatest diversity of bacteria: BHI-YEc, the non-selective Brain Heart Infusion medium (BD, Franklin Lakes, NJ, USA) supplemented with 1% yeast extract (Biokar, Pantin, France) and 0.05% L-cysteine hydrochloride (Sigma-Aldrich, St. Quentin Fallavier, France) to promote the growth of all HM bacteria; the Columbia II medium containing 5% sheep blood (BD, Le Pont de Claix, France) (BA), which promotes the growth of fastidious microorganisms; the Yeast extract Casitone and Fatty Acids (YCFA), the Peptone Yeast Glucose (PYG) modified medium, and the Wilkins Chalgren (BD, Franklin Lakes, NJ, USA) medium supplemented with 0.05% L-cysteine-hydrochloride (WCc), these three media favouring the growth of anaerobic bacteria; and the Man Rogosa Sharp (BD, Franklin Lakes, NJ, USA) medium supplemented with 0.05% L-cysteine-hydrochloride (MRSc) to favour the growth of Lactobacilli. All media were incubated at 37°C in anaerobic jars for 1 to 3 days, except BHI-YEc which was incubated in both aerobic and anaerobic conditions. All the colonies on the surface of each medium were collected by scrapping the plate, resuspended in 500 µL of 0.9% NaCl solution, pooled and centrifuged (10,000g, 5 min, 4°C) and the pellet was stored at-80°C until microbiota analysis by metabarcoding in order to determine the complete CM.

### Raw and cultivable milk microbiota analysis

Microbiota analysis by metabarcoding was performed on RM and CM. Raw milk samples (3 mL) were thawed on ice, mixed with 1 mL of sodium citrate (1 M, pH 7.5) and centrifuged (18,000g, 20 min, 4°C). After washing with 1 mL of sodium citrate (20 g/L, pH 7.5) and centrifugation (18,000g, 15 min, 4°C), the pellet was resuspended in 100 μL TE buffer (10 mM TRIS–HCl (pH8), 2 mM EDTA). For CM, frozen pellets were resuspended directly in 100 μL TE buffer.

DNA extraction and PCR amplification were performed exactly as described by Mariadassou et al, using the universal primers S-D-Bact-0341-b-S-17 and S-D-Bact-0785-a-A-21 targeting the V3-4 region of the gene encoding 16S rRNA ^59^. Amplicon sequencing on the Illumina MiSeq PE250 platform (Illumina Inc., San Diego, CA, USA) from Genome Quebec (Montreal, Canada) was performed as previously described ^59^. Negative controls that underwent all steps from extraction to sequencing but without bacterial suspension were included for each set of extractions, resulting in 6 negative controls.

Sequence library analysis was performed using the FROGS pipeline hosted on the INRAE MIGALE bioinformatics platform essentially as previously described ^60^. Briefly, the preprocessing, clustering and chimera removal steps were performed using the FROGS pipeline (Galaxy version 3.2.3+galaxy2) as previously described ^59^. The FROGS clustering step was performed using Swarm with an aggregation distance of 1, allowing for one mismatch between clustered sequences ^61^. The data were filtered using the FROGS OTU filter tool, to retain OTUs with a minimum proportion of 0.00005 in the complete dataset. Affiliation was then performed with the FROGS affiliation OTU tool based on Blastn+ using the 16S SILVA 138.1 database ^62,63^. The median number of sequences was 53 465 sequences per sample. Negative controls reached a median number of 25 sequences per sample, which was considered negligible compared to milk samples. Negative controls were therefore not included in the analysis. Finally, a rooted phylogenetic tree was created with FastTree and Phangorn R package implemented on FROGS pipeline.

Additional filters were considered to determine the number of taxa (from phylum to OTU) in each sample and prevent its overestimation. Only taxa with a total number of reads ≥ 8 in a given sample were included in the number of taxa in this sample (this corresponded to a minimum proportion of 0.00005 in the sample with the highest number of reads). Likewise, only taxa that were present at least in one CM or one RM sample with a number of reads ≥ 8 were considered in the total number of taxa in CM and RM respectively.

### Digestion of selected strains in an infant formula matrix

Six strains isolated from the same HM samples and belonging to prevalent genera of HM microbiota were used, namely *Bifidobacterium breve* CIRM BIA 2845, *Streptococcus salivarius* CIRM BIA 2846, *Cutibacterium acnes* CIRM BIA 2849, *Staphylococcus epidermidis* CIRM BP 1633, *Lactobacillus gasseri* CIRM BIA 2841 and *Enterococcus faecalis* CIRM BIA 2835, (hereafter referred to without the strain number) ^24^.

Strains were cultivated at 37°C under static condition for 1 to 3 days, in MRSc for *B. breve* and *L. gasseri* in anaerobic jar, in PYG for *C. acnes* in anaerobic jar, in Tryptic soy broth (Biokar, Pantin, France) supplemented with 1% yeast extract (TSE) for *E. faecalis* and *S. salivarius*, and in BHI-YEc for *S. epidermidis.* The bacterial concentration of the culture was then determined by measuring the optical density at 600 nm (VWR V-1200 Visible Spectrophotometer, VWR, Rosny-sous-Bois, France) and using a previously established OD/CFU relationship. Bacteria were centrifuged (6800 g, 5 min, RT), washed in HBSS 1X (Dutscher, Bernolsheim, France), centrifuged (6800 g, 5 min, RT) and resuspended in RPMI 1640 containing glutamine but without antibiotics or SVF, at a concentration of 1×10^9^ CFU/mL. Part of this suspension was used to assess the effect of the strain on THP1 cells before digestion (condition “B”).

The remaining suspension was centrifuged (6800 g, 5 min, RT) and resuspended in an infant formula (IF) (composition for 100 mL as follows: 3.43 g proteins, 8.42 g lipids, 6.21g carbohydrates, 1.34 g ashes and vitamins) at a bacterial concentration of 3.8×10^7^ CFU/mL before digestion using an *in vitro* model in static conditions at the full-term infant stage ^29^. This method mimics *in vitro* the enzymatic and physicochemical parameters of gastro-intestinal digestion of breast milk in a full-term infant (28 days) weighing approximately 4.25 kg. Two phases were simulated. For the gastric phase, IFs containing bacteria were digested for 1 h at pH 5.3, with simulated gastric fluid and porcine pepsin. Lipase was not used in this model due to limited commercial availability. These digesta were then digested for 1 h at pH 6.6 in a simulated intestinal fluid containing bovine bile and porcine pancreatin (lyophilized pancreatic juice containing trypsin, chymotrypsin and lipase) to mimic the intestinal digestion ^29^. Digestion was performed in three independent biological replicates, each with two technical replicates.

Bacterial cultivability was determined after the gastric and intestinal digestions (conditions “G” and “I” respectively) directly on the digesta using the media described above and the micromethod as previously described ^64^.

The remaining digesta was then centrifuged (6800 g, 10 min, RT), washed once with HBSS (6800 g, 10 min, RT) and resuspended in RPMI 1640 containing glutamine but without antibiotics or SVF. The theoretical concentration of this bacterial suspension was 1×10^9^ CFU/mL, with no effect of digestion on the cultivability of the strain. A control corresponding to the bacteria-free IF was also digested, washed (under the same conditions as the bacterial pellet) and used for interaction with THP1 cells to assess the immunomodulatory effect of the digested matrix (condition “IF-I”; see below).

### Immunomodulatory properties of bacteria on THP1 cells differentiated into macrophages

The immunomodulatory potential of bacteria was assessed on the monocyte THP1 cell line (ATCC TIB-202). Cells were grown in RPMI1640 (Merck Sigma, Darmstadt, Germany) containing glutamine, supplemented with 10% SVF (decomplemented by heating for 30 min at 56°C) (Biowest, Riverside, United States), 100 IU/mL penicillin, and 100 μg/mL streptomycin (Merck Sigma) (hereafter referred to as RPMIc). Cells were grown at 37°C in a 5% CO2 water-saturated atmosphere in 25 or 75 cm^2^ flasks (Corning Inc, Corning, NY, USA) and diluted to 3×10^5^ cells/mL in RPMIc every 3 days. THP1 cells were differentiated into macrophages two days before use. Briefly, 5×10^5^ cells/well were seeded in 48-well plates and incubated in RPMIc with 200nM PMA (Phorbol 12-myristate 13-acetate, Merck Sigma, Saint Quentin Fallavier, France). After 24 h, cells differentiated into macrophages were adherent. They were washed 3 times with HBSS (Merck Sigma, Saint Quentin Fallavier, France) before adding 500 µl RPMIc for a further 24 h incubation before the interaction with bacteria.

The bacterial suspension at 1×10^9^ CFU/mL, either before digestion or after recovery from intestinal digestion, was diluted at 2×10^7^ CFU/mL in RPMIc. THP1 cells (5×10^5^ cells/well) were stimulated at a multiplicity of infection (MOI) of 10:1 bacteria per cell, for 24 h in RPMIc at 37°C in a 5% CO_2_ water saturated atmosphere. The supernatant was centrifuged (8000 g, 5 min, 4°C) and, after addition of anti-protease cocktail 1X (SigmaFast, Merck Sigma, Saint Quentin Fallavier, France), stored at-20°C until cytokine analysis by ELISA assay.

IL-10 and TNF-α were then determined by ELISA (Human IL-10 ELISA Set, Cat. No. 555157; Human TNF-α ELISA Set, Cat. No. 555212; TMB Substrate Reagent Set, Cat. No. 555214) according to the manufacturer’s recommendations. Washes were performed using a biotek 50TS8V 8-channel microplate washer (Agilent Technologies, Les Ulis, France), and absorbance at 450 nm was measured by the biotek 800TS microplate reader (Agilent Technologies, Les Ulis, France). IL-10 and TNF-α production (pg/mL) were corrected by the production in the absence of bacterial stimulation (C, control) of the corresponding replicate.

## Statistical analysis

Statistical analyses on microbiota were performed using R 4.3.0 and specialized packages: phyloseq (v. 1.34), DESeq2 (v 1.30.1) and custom scripts ^59,65–67^. Statistical analyses were performed considering separately raw milk (RM) and cultivable (CM) milk microbiota or both types of samples (whole dataset). Several factors were considered for the analysis, including parity, mother’s age and body mass index (BMI) and maternal supplementation with iron and/or with vitamins and trace elements either during gestation (iron_G and vit_G) or during breastfeeding (iron_BF and vit_BF) (Supplementary Table S1). Additional metadata are shown in Supplementary Table S1 but were not considered due to a limited number of individuals per modality (e.g. diet) or redundancy with other factors (height and weight with BMI).

The analysis on α-and β-diversity indices was performed on data rarefied to the same depth. Observed, Shannon and Inversed Simpson indices were used to represent the α-diversity. The impact of the different factors on each index was first assessed using a multivariate analysis of variance (ANOVA) (P-value < 0.05) considering the following factors: parity, mother’s age, BMI, iron_G, vit_G, iron_BF, vit_BF and sample type (on the whole dataset only). As mother’s age, BMI, vit_G and iron_BF were not significant (p > 0.05), these factors were removed from the model. Beta diversity analyses were performed using the UniFrac, weighted UniFrac, Bray-Curtis and Jaccard distances. MultiDimensional Scaling (MDS) was performed on the weighted UniFrac and Bray-Curtis distance matrix to represent the samples on the principal plane. The effect of the different factors on β-diversity was assessed using multivariate analysis of variance (PERMANOVA), as implemented in the adonis2 function from the vegan package, and considering the same factors as for α-diversity. As no impact on the β-diversity was observed with BMI, mother’s age, vit_G, vit_BF and iron_BF (p > 0.05), these factors were removed from the model.

Finally, differences in phyla, classes, orders, families, genera and OTUs were assessed with pairwise comparisons for the different factors affecting α-and β-diversities. This differential abundance analysis was performed using the DESeq2 R package implemented in the Easy16S R-shiny interface hosted on the Shiny Migale platform ^66^ after aggregation to the desired taxonomic rank (phyloseq tax_glom function).

Regarding the effect of digestion on bacterial cultivability and immunomodulatory properties, statistical analysis was performed using R software (version 4.3.0, ^67^). Differences between groups were examined using a one-way analysis of variance (ANOVA), followed by the Tukey significant difference test to compare group means. Differences were considered statistically significant at p < 0.05.

## Data availability

Data files related to metagenomic analysis are available at the Sequence Read Archive of the National Center for Biotechnology Information under the accession number PRJNA1184131.

## Ethics declaration

The HM sampling protocol was approved by the Institutional Review Board of the Poitiers Hospital (n°20.05.27.67526). Informed consent was obtained from all participants.

## Acknowledgments

We warmly thank all volunteer mothers for their breastmilk samples and all the paediatric nurses of the maternity department of Rennes University Hospital, for their involvement in the project. We thank all the members from the sequencing platform of Genome Quebec for their help and all their efforts in the realization of this project, with special thanks to, Mrs Julie Boudeau, Mrs Hao Fan Yam and Mr Frederick Robidoux. We are grateful to the INRAE MIGALE bioinformatics facility (MIGALE, INRAE, 2020. Migale bioinformatics Facility, doi: 10.15454/1.5572390655343293E12) for providing computing and storage resources.

## Authors contributions

SE, ILHL and YLL conceived the project that led to the submission of this work. CLB, SE, ILHL, AB and AD designed the experiment. NJ, VC, FV and LR collected human milk. CLB, AM, LR, MFC, OM and NJ performed experiments and acquired data. CLB, AM, ILHL and SE interpreted the data. CLB, ILHL and SE drafted the manuscript and all authors revised and approved it.

## Data availability statement

All data generated or analysed during this study are included in this published article and its supplementary information files.

**Additional Information (including a Competing Interests Statement) Supplementary Figure S1**. Prevalent genera in the cultivable milk microbiota

**Supplementary Figure S2.** Impact of different factors on the alpha diversity of raw (**a,c**) and cultivable (**b,d**) milk microbiota.

**Supplementary Table S1**. Human milk (HM) samples and associated metadata. For each sample, microbiota was determined on raw milk and on the cultivable fraction

**Supplementary Table S2**. Abundance table in each sample (either raw milk microbiota (RM) or cultivable milk microbiota (CM), aggregated at different taxonomic levels, from phylum to OTU

**Supplementary Table S3**. Differentially abundant taxa in raw milk microbiota (RM) with regard to different factors

**Supplementary Table S4**. Differentially abundant taxa in cultivable milk microbiota (CM) with regard to different factors

## References

1. Jost, T., Lacroix, C., Braegger, C. & Chassard, C. Assessment of bacterial diversity in breast milk using culture-dependent and culture-independent approaches. Br.J.Nutr. 110, 1253–1262 (2013).

2. Togo, A. et al. Repertoire of Human Breast and Milk Microbiota: A Systematic Review. Future Microbiol. 14, 623–641 (2019).

3. Fernández, L., Pannaraj, P. S., Rautava, S. & Rodríguez, J. M. The Microbiota of the Human Mammary Ecosystem. Front. Cell. Infect. Microbiol. 10, (2020).

4. Oikonomou, G. et al. Milk Microbiota: What Are We Exactly Talking About? Front. Microbiol. 11, 60 (2020).

5. Notarbartolo, V., Giuffrè, M., Montante, C., Corsello, G. & Carta, M. Composition of Human Breast Milk Microbiota and Its Role in Children’s Health. Pediatr. Gastroenterol. Hepatol. Nutr. 25, 194–210 (2022).

6. Sun, Q. et al. Effects of Maternal Diet on Infant Health: A Review Based on Entero-Mammary Pathway of Intestinal Microbiota. Mol. Nutr. Food Res. 2400077 (2024) doi:10.1002/mnfr.202400077.

7. Zimmermann, P. & Curtis, N. Breast milk microbiota: A review of the factors that influence composition. J. Infect. 81, 17–47 (2020).

8. Li, C. et al. CcpA mediates proline auxotrophy and is required for the pathogenesis of Staphylococcus aureus infections. J. Bacteriol. (2010) doi:10.1128/JB.00237-10.

9. Ding, M. et al. Geographical location specific composition of cultured microbiota and Lactobacillus occurrence in human breast milk in China. Food Funct. 10, 554–564 (2019).

10. Schwab, C., Voney, E., Ramirez Garcia, A., Vischer, M. & Lacroix, C. Characterization of the Cultivable Microbiota in Fresh and Stored Mature Human Breast Milk. Front. Microbiol. 10, 2666 (2019).

11. Treven, P. et al. Evaluation of Human Milk Microbiota by 16S rRNA Gene Next-Generation Sequencing (NGS) and Cultivation/MALDI-TOF Mass Spectrometry Identification. Front. Microbiol. 10, (2019).

12. Boudry, G. et al. The Relationship Between Breast Milk Components and the Infant Gut Microbiota. Front. Nutr. 8, 629740 (2021).

13. Selma-Royo, M. et al. Human milk microbiota: what did we learn in the last 20 years? Microbiome Res. Rep. 1, 19 (2022).

14. Williams, J. E. et al. Strong Multivariate Relations Exist Among Milk, Oral, and Fecal Microbiomes in Mother-Infant Dyads During the First Six Months Postpartum. J. Nutr. 149, 902–914 (2019).

15. Fehr, K. et al. Breastmilk Feeding Practices Are Associated with the Co-Occurrence of Bacteria in Mothers’ Milk and the Infant Gut: the CHILD Cohort Study. Cell Host Microbe 28, 285–297.e4 (2020).

16. Laursen, M. F. et al. Maternal milk microbiota and oligosaccharides contribute to the infant gut microbiota assembly. ISME Commun. 1, 1–13 (2021).

17. Asnicar, F. et al. Studying Vertical Microbiome Transmission from Mothers to Infants by Strain-Level Metagenomic Profiling. mSystems 2, e00164–16 (2017).

18. Murphy, K. et al. The Composition of Human Milk and Infant Faecal Microbiota Over the First Three Months of Life: A Pilot Study. Sci. Rep. 7, 40597 (2017).

19. Liu, H. et al. Maternal milk and fecal microbes guide the spatiotemporal development of mucosa-associated microbiota and barrier function in the porcine neonatal gut. BMC Biol. 17, 106 (2019).

20. Díaz-Ropero, M. P. et al. Two Lactobacillus strains, isolated from breast milk, differently modulate the immune response. J. Appl. Microbiol. 102, 337–343 (2007).

21. Jeon, S. G. et al. Probiotic Bifidobacterium breve induces IL-10-producing Tr1 cells in the colon. PLoS Pathog. 8, e1002714 (2012).

22. Al-Sadi, R. et al. Bifidobacterium bifidum Enhances the Intestinal Epithelial Tight Junction Barrier and Protects against Intestinal Inflammation by Targeting the Toll-like Receptor-2 Pathway in an NF-κB-Independent Manner. Int. J. Mol. Sci. 22, 8070 (2021).

23. Li, S. et al. Gut Microbiota and Immune Modulatory Properties of Human Breast Milk Streptococcus salivarius and S. parasanguinis Strains. Front. Nutr. 9, (2022).

24. Le Bras, C. et al. Two human milk-like synthetic bacterial communities displayed contrasted impacts on barrier and immune responses in an intestinal quadricellular model. ISME Commun. ycad019 (2024) doi:10.1093/ismeco/ycad019.

25. Bezkorovainy, A. Probiotics: determinants of survival and growth in the gut123. Am. J. Clin. Nutr. 73, 399s–405s (2001).

26. Lo Curto, A., et al. Survival of probiotic lactobacilli in the upper gastrointestinal tract using an in vitro gastric model of digestion. Food Microbiol. 28, 1359–1366 (2011).

27. Pitino, I. et al. Survival of Lactobacillus rhamnosus strains inoculated in cheese matrix during simulated human digestion. Food Microbiol. 31, 57–63 (2012).

28. Kowalczyk, M., Znamirowska-Piotrowska, A., Buniowska-Olejnik, M. & Pawlos, M. Sheep Milk Symbiotic Ice Cream: Effect of Inulin and Apple Fiber on the Survival of Five Probiotic Bacterial Strains during Simulated In Vitro Digestion Conditions. Nutrients 14, 4454 (2022).

29. Ménard, O. et al. A first step towards a consensus static in vitro model for simulating full-term infant digestion. Food Chem. 240, 338–345 (2018).

30. Blanchet, F. et al. Heat inactivation partially preserved barrier and immunomodulatory effects of Lactobacillus gasseri LA806 in an in vitro model of bovine mastitis. Benef. Microbes 1–12 (2021) doi:10.3920/BM2020.0146.

31. Chang, Y. et al. Optimization of Culturomics Strategy in Human Fecal Samples. Front. Microbiol. 10, (2019).

32. Wang, F. et al. An optimized culturomics strategy for isolation of human milk microbiota. Front. Microbiol. 15, (2024).

33. De Andrés, J. et al. Physiological Translocation of Lactic Acid Bacteria during Pregnancy Contributes to the Composition of the Milk Microbiota in Mice. Nutrients 10, 14 (2018).

34. Lagier, J.-C. et al. Culturing the human microbiota and culturomics. Nat. Rev. Microbiol. 16, 540– 550 (2018).

35. Melchior, S. et al. Effect of the formulation and structure of monoglyceride-based gels on the viability of probiotic Lactobacillus rhamnosus upon in vitro digestion. Food Funct. 12, 351–361 (2021).

36. Liu, W. et al. Characterization of potentially probiotic lactic acid bacteria and bifidobacteria isolated from human colostrum. J. Dairy Sci. (2020) doi:10.3168/jds.2019-17602.

37. Papadimitriou, K. et al. Stress Physiology of Lactic Acid Bacteria. Microbiol. Mol. Biol. Rev. MMBR 80, 837–890 (2016).

38. Horáčková, Š., Plocková, M. & Demnerová, K. Importance of microbial defence systems to bile salts and mechanisms of serum cholesterol reduction. Biotechnol. Adv. 36, 682–690 (2018).

39. Gao, X. et al. Lactobacillus, Bifidobacterium and Lactococcus response to environmental stress: Mechanisms and application of cross-protection to improve resistance against freeze-drying. J. Appl. Microbiol. 132, 802–821 (2022).

40. Barnes, A. M. T. et al. Enterococcus faecalis readily colonizes the entire gastrointestinal tract and forms biofilms in a germ-free mouse model. Virulence 8, 282–296 (2017).

41. Rabah, H. et al. Cheese matrix protects the immunomodulatory surface protein SlpB of Propionibacterium freudenreichii during in vitro digestion. Food Res. Int. Ott. Ont 106, 712–721 (2018).

42. Adouard, N. et al. Survival of cheese-ripening microorganisms in a dynamic simulator of the gastrointestinal tract. Food Microbiol. 53, 30–40 (2016).

43. Zhou, C. & Fey, P. D. The Acid Response Network of Staphylococcus aureus. Curr. Opin. Microbiol. 55, 67–73 (2020).

44. Alsultan, A., Walton, G., Andrews, S. C. & Clarke, S. R. Staphylococcus aureus FadB is a dehydrogenase that mediates cholate resistance and survival under human colonic conditions. Microbiology 169, 001314 (2023).

45. Chauvet, L. et al. Protein ingredient quality of infant formulas impacts their structure and kinetics of proteolysis under *in vitro* dynamic digestion. Food Res. Int. 169, 112883 (2023).

46. Phillips-Farfán, B. et al. Microbiota Signals during the Neonatal Period Forge Life-Long Immune Responses. Int. J. Mol. Sci. 22, 8162 (2021).

47. Gavzy, S. J. et al. Bifidobacterium mechanisms of immune modulation and tolerance. Gut Microbes 15, 2291164 (2023).

48. Yin, R. et al. Immunogenic molecules associated with gut bacterial cell walls: chemical structures, immune-modulating functions, and mechanisms. Protein Cell 14, 776–785 (2023).

49. Moossavi, S. et al. Composition and Variation of the Human Milk Microbiota Are Influenced by Maternal and Early-Life Factors. Cell Host Microbe 25, 324–335.e4 (2019).

50. Kordy, K. et al. Contributions to human breast milk microbiome and enteromammary transfer of Bifidobacterium breve. PLoS ONE 15, (2020).

51. Cortes-Macías, E. et al. Maternal Diet Shapes the Breast Milk Microbiota Composition and Diversity: Impact of Mode of Delivery and Antibiotic Exposure. J. Nutr. 151, 330–340 (2021).

52. Williams, J. E. et al. Human Milk Microbial Community Structure Is Relatively Stable and Related to Variations in Macronutrient and Micronutrient Intakes in Healthy Lactating Women. J. Nutr. 147, 1739–1748 (2017).

53. Moro, G. E. et al. Adherence to the Traditional Mediterranean Diet and Human Milk Composition: Rationale, Design, and Subject Characteristics of the MEDIDIET Study. Front. Pediatr. 7, (2019).

54. Babakobi, M. D. et al. Effect of Maternal Diet and Milk Lipid Composition on the Infant Gut and Maternal Milk Microbiomes. Nutrients 12, 2539 (2020).

55. Padilha, M. et al. The Human Milk Microbiota is Modulated by Maternal Diet. Microorganisms 7, 502 (2019).

56. Finlayson-Trick, E. C., Fischer, J. A., Goldfarb, D. M. & Karakochuk, C. D. The Effects of Iron Supplementation and Fortification on the Gut Microbiota: A Review. Gastrointest. Disord. 2, 327–340 (2020).

57. Pereira, D. I. A. et al. Dietary iron depletion at weaning imprints low microbiome diversity and this is not recovered with oral nano Fe(III). MicrobiologyOpen 4, 12–27 (2015).

58. Nitert, M. D. et al. Iron supplementation has minor effects on gut microbiota composition in overweight and obese women in early pregnancy. Br. J. Nutr. 120, 283–289 (2018).

59. Mariadassou, M. et al. Microbiota members from body sites of dairy cows are largely shared within individual hosts throughout lactation but sharing is limited in the herd. *Anim*. Microbiome 5, 32 (2023).

60. Goetz, C. et al. Post-milking application of a Lacticaseibacillus paracasei strain impacts bovine teat microbiota while preserving the mammary gland physiology and immunity. Benef. Microbes 15, 275–291 (2024).

61. Mahé, F., Rognes, T., Quince, C., de Vargas, C. & Dunthorn, M. Swarm: robust and fast clustering method for amplicon-based studies. PeerJ 2, e593 (2014).

62. Camacho, C. et al. BLAST+: architecture and applications. BMC Bioinformatics 10, 421 (2009).

63. Quast, C. et al. The SILVA ribosomal RNA gene database project: improved data processing and web-based tools. Nucleic Acids Res 41, D590–D596 (2013).

64. Baron, F. et al. Rapid and cost-effective method for microorganism enumeration based on miniaturization of the conventional plate-counting technique. Lait 86, 251–257 (2006).

65. McMurdie, P. J. & Holmes, S. phyloseq: an R package for reproducible interactive analysis and graphics of microbiome census data. PloS One 8, e61217 (2013).

66. Love, M. I., Huber, W. & Anders, S. Moderated estimation of fold change and dispersion for RNA-seq data with DESeq2. Genome Biol. 15, 550 (2014).

67. R development Core Team. R: A Language and Environment for Statistical computing (url:http://www.r-project.org). R Foundation for Statistical Computing (2023).

