## Supplementary figure S1 for "New insights into the cultivability of human milk microbiota from ingestion to digestion and implications for its immunomodulatory properties"

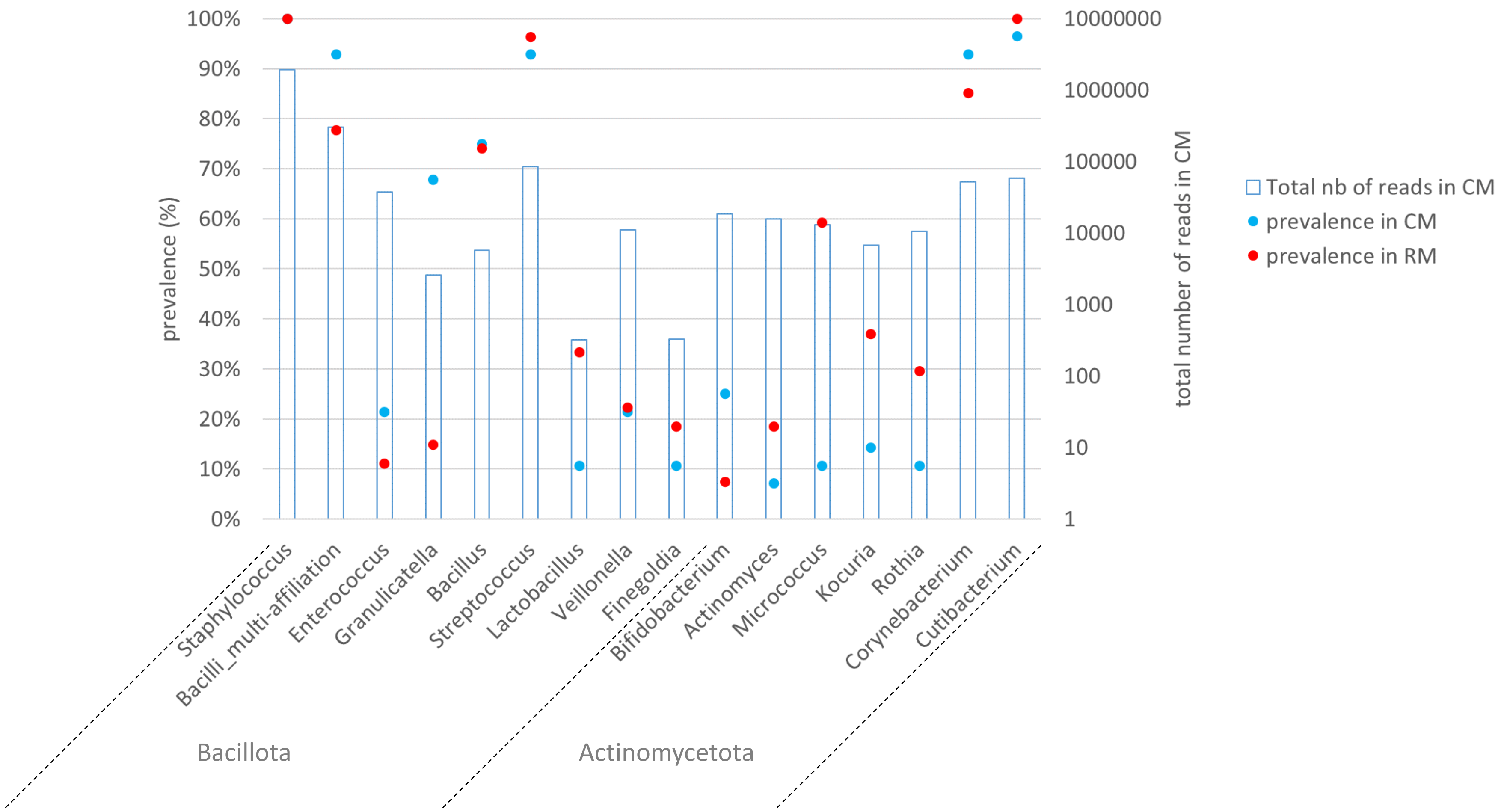


**Supplementary Figure S1**. Prevalent genera in the cultivable milk microbiota

Genera with prevalence in the cultivable milk microbiota ( ● ) higher than 7% (corresponding to a minimum of 2 HM samples) and with a total relative abundance in CM higher than 0.00005 (bar chart) are presented, as well as the prevalence of these genera in the raw milk microbiota (RM) ( ● ).
