## Supplementary figure S2 for "New insights into the cultivability of human milk microbiota from ingestion to digestion and implications for its immunomodulatory properties"

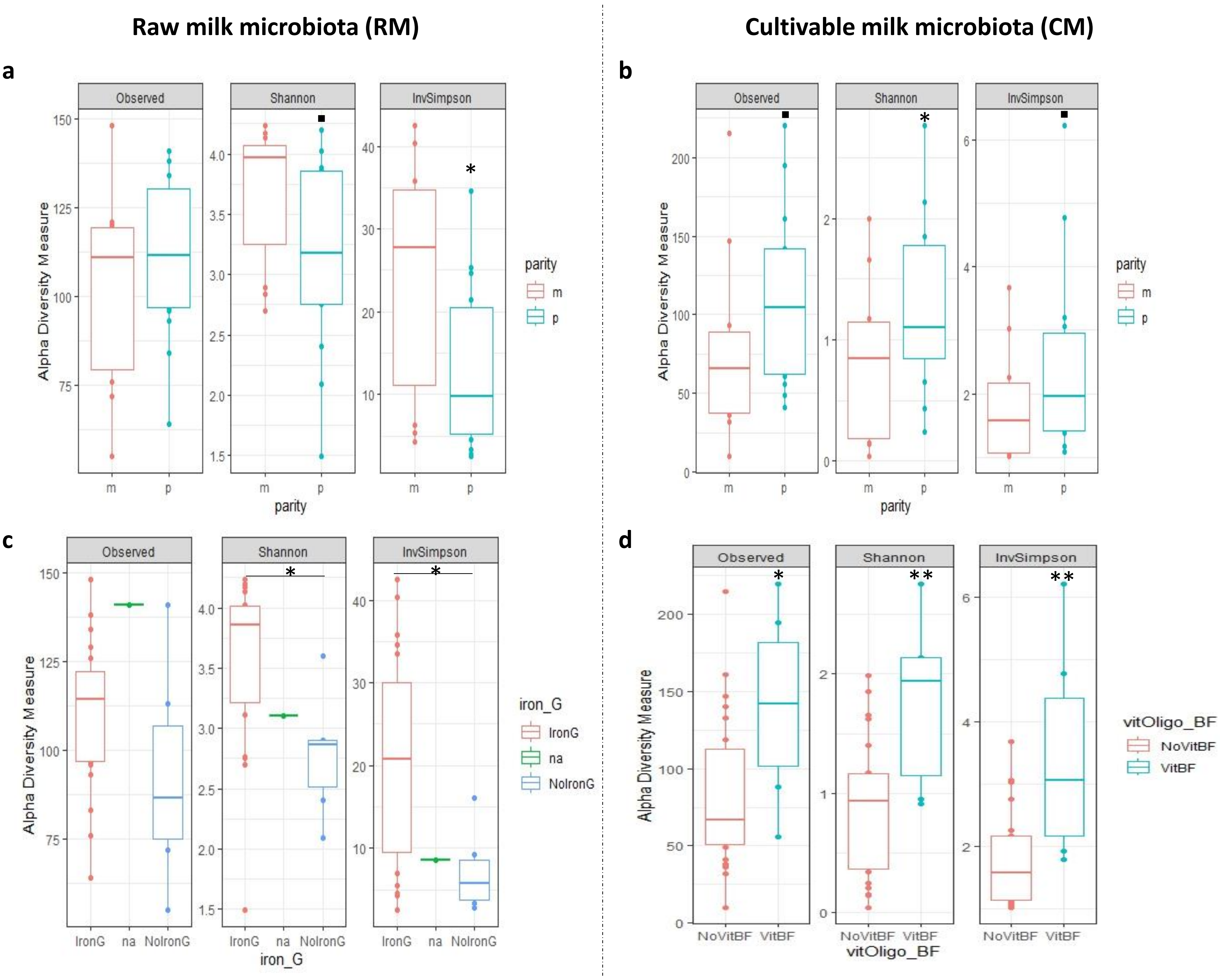


**Supplementary Figure S2.** Impact of different factors on the alpha diversity of raw (**a,c**) and cultivable (**b,d**) milk microbiota. Three alpha diversity indexes were used: Observed richness, Shannon and Inverse Simpson; (**a,b**) Parity : either primiparous (p) or multiparous (m); (**c**) Iron_G: mothers supplemented or not by iron during gestation; (**d**) vitOligo_BF: mothers supplemented or not by vitamins during breastfeeding ; na: missing metadata. *pval<0.05, ** pval<0.01, ▪ P value < 0.1
